# The selection of antibiotic- and bacteriophage-resistant *Pseudomonas aeruginosa* is prevented by their combination

**DOI:** 10.1101/2022.05.25.493369

**Authors:** Aude A Ferran, Marlène Z. Lacroix, Ophélie Gourbeyre, Alicia Huesca, Baptiste Gaborieau, Laurent Debarbieux, Alain Bousquet-Mélou

## Abstract

**Objectives:** Bacteria developing resistance compromise the efficacy of antibiotics or bacteriophages (phages). We tested the association of these two antibacterials to circumvent resistance.

**Methods:** With the Hollow Fiber Infection Model (HFIM), we mimicked the concentration profile of ciprofloxacin in the lungs of patients treated orally for *Pseudomonas aeruginosa* infections and independently, mimicked a single inhaled administration of phages (one or two phages).

**Results:** Each treatment selects for antibiotic-or phage-resistant clones in less than 30 h. By contrast, no bacteria were recovered from the HFIM at 72 h when ciprofloxacin was started 4 h post-phage administration, even when increasing the initial bacterial concentration by a 1000 fold.

**Conclusion:** The combination of phages with antibiotics used according to clinical regimens prevents the growth of resistant clones, providing opportunities to downscale the use of multiple antibiotics.

## Introduction

*Pseudomonas aeruginosa* is an opportunistic pathogen, naturally resistant to many antibiotics. Moreover, repeated antibiotics treatments administered to patients with chronic airways infections, such as cystic fibrosis (CF) patients, have led this bacterium to acquire additional drug-resistance [1,2]. Acute exacerbations are treated by either intravenous (beta-lactams and aminoglycosides), oral (ciprofloxacin), or inhaled (tobramycin and colistimethate sodium) administrations of antibiotics [2,3]. The recent TORPEDO-CF study concluded that IV or oral antibiotics administration could be equivalent [2].

In *P. aeruginosa* resistance to fluoroquinolones, such as ciprofloxacin, involves mutations in *gyrA* or in genes regulating the expression of the MexEF-OprN efflux pump [4,5]. The increase of the MIC is often modest for the first-step mutants, qualified as “less-susceptible” [5], but is sufficient to favor their growth within the Mutant Selection Window (MSW) [6]. Next, multiple mutations lead to a higher MIC, clinical resistance [4], and ultimately require ciprofloxacin to be associated with other antibiotics, upscaling drug use [7,8]. Interestingly, the gradual increase of MIC is reproduced *in vitro* by mimicking the clinical regimens of ciprofloxacin, providing a mean to study the efficacy of combined treatments [9–11].

Bacteriophages (phages) are antibacterial viruses. Recently, an increasing number of successful compassionate treatments in both Europe and the USA have confirmed the therapeutic potential of phages, which has a long history of human use [12,13]. Phages have the peculiar capacity to self-amplify at the site of infection, increasing their density locally, at the expense of bacteria [14]. Nevertheless, as for antibiotics, bacteria have developed several ways to resist to phages [15]. However, since the molecular mechanisms involved in drug and phage resistance do not overlap, their association in cocktails or with antibiotics has been proposed to improve the efficacy of treatments [16,17].

By using the Hollow Fiber Infection Model (HFIM, Fig S1, S2) inoculated with *P. aeruginosa*, we simulated the pharmacokinetics of ciprofloxacin in the human lungs for 72 h and evaluated the antibacterial efficacy of its combination with phages administered to mimic a single local inhaled treatment. We show that the combination of phages with a simultaneous or delayed administration of ciprofloxacin leads to a stronger reduction of *P. aeruginosa* in the HFIM than individual treatments, preventing the selection of both phage and ciprofloxacin resistant clones. This reduction reached the limit of detection with the delayed combinations, suggesting that coupling phages and antibiotics could downscale antibiotics consumption in clinics.

## Material and methods

### Bacterial strain

The *Pseudomonas aeruginosa* strain K (PAK) with a MIC of ciprofloxacin of 0.064 µg/mL was used for all the experiments.

### Phages and ciprofloxacin

Phage PAK_P1, a virulent *Myoviridae*, was isolated using strain PAK[18]. Phage LUZ19v is a variant of phage LUZ19, a virulent *Podoviridae* isolated on strain PAO1 [21], isolated following serial passages on strain PAK. The efficiency of plating (EOP) of LUZ19 and LUZ19v on strain PAK is 0.2 and 1, respectively, compared to the EOP on strain PAO1

Both phages were amplified in liquid lysogeny broth. Lysates were filtered-sterilized at 0.2µm, and stored at 4°C until use. Phage titrations (serial dilutions) were spotted on tryptic soy agar (TSA) supplemented with magnesium sulfate (10 g/L) and activated charcoal (10 g/L) covered by a lawn of strain PAK made with 10^6^ CFU.

Stocks of ciprofloxacin (Sigma Aldrich) were stored at -20°C for less than 1 month and thawed only once.

### Minimal Inhibitory Concentration (MIC) determination

The MIC of ciprofloxacin for the strain PAK was determined in triplicate by broth microdilution in cation-adjusted Mueller Hinton broth (MHB), according to the CLSI reference methods. Frozen samples of bacteria collected at the end (72 h) of each HFIM experiment were thawed and plated on MH agar overnight. Several colonies were sampled and the bacterial density was adjusted to 5×10^5^ CFU/mL before MIC determination.

### Hollow Fiber Infection Model (HFIM)

The HFIM includes a cartridge (C2011 polysulfone cartridge, FiberCell Systems, Inc., Frederick, MD, USA) with capillaries composed of a semipermeable polysulfone membrane. The pore size of the capillaries (42 kDa) allows equilibration of ciprofloxacin, which can freely circulate between the intracapillary and extracapillary spaces while the bacteria and the phages are trapped in the extracapillary space of the cartridge (Fig. S1 and S2).

In this study, 20 mL of a suspension containing 5.5 log_10_ CFU/mL (standard inoculum) or 8.5 CFU/mL (high inoculum) of *P. aeruginosa* were inoculated into the extracapillary space of each cartridge and incubated at 37°C in MHB for 1 h. Ciprofloxacin was added to the central compartment to obtain the desired maximum concentration (C_max_) of 1.5 µg/mL and was continuously diluted with MHB to mimic an elimination half-life of 4 h [19]. A mean inoculum of 7.5 log_10_ PFU/mL (8.8 log_10_ PFU *in toto*) of either one or two phages with equal amounts of each phage was added once into the extracapillary space. Treatments with ciprofloxacin or phages or both simultaneously were started 1 h after the inoculation of bacteria in the HFIM. When testing the delayed combination, ciprofloxacin was added 4 h after the phages. All the experiments (except the untreated control and single phage combined to ciprofloxacin) were performed in duplicate.

### Bacteria and phages quantification

Samples of 1 mL were collected from the extracapillary space to count the bacteria and phages at 0 (before the addition of phages or ciprofloxacin), 0.25, 0.5, 1, 2, 4, 6, 8, 24, 26, 29, 32, 48, 50, 53, 56, and 72 h. After centrifugation at 3000 g for 10 min, supernatants were recovered to count phages and pellets were resuspended in 1 mL of NaCl 0.9% to count bacteria. The bacteria and phage suspensions were serially diluted (10X), and spotted (10 µL) in triplicate on either TSA (to count bacteria) or on TSA covered with strain PAK (to count phages) and incubated overnight at 37°C. The limit of detection (LOD) was 1.5 log_10_ CFU/mL for bacteria and 1.5 log_10_PFU/mL for phages.

### Monitoring of ciprofloxacin and phage resistance

Twice a day the bacteria sampled from the HFIM were counted on agar plates containing 0.5 µg/mL of ciprofloxacin, corresponding to 8-fold the MIC of t strain PAK. The proportion of less-susceptible bacteria was calculated as the ratio of colonies on drug-supplemented agar (MIC 8X) divided by colonies on drug-free agar.

Bacteria sampled at 72 h were stored in 30% v/v glycerol at -80°C before phage resistance analysis. Bacteria were thawed and immediately incubated with or without phages (10^8^ PFU) for one hour before plating on agar with or without a single pre-absorbed phage (10^8^ PFU/plate). The successive incubations in broth and on agar were made with the same phage (LUZ19v or PAK_P1). The frequency of resistance to each phage was calculated by the ratio of colonies growing in the presence of phages over those growing in the absence of phages.

### Ciprofloxacin quantification

Ciprofloxacin quantification was performed in samples withdrawn from the central compartment and from the extracapillary space of the cartridge according to a standard protocol (Supplementary file).

## Results

### Pharmacokinetic analysis of ciprofloxacin in the HFIM

We simulated in the HFIM inoculated with *P. aeruginosa* strain PAK the concentration profile of ciprofloxacin during 72 h corresponding to the administration of 500 mg twice daily in patients, using a C_max_ at 1.5 µg/mL and a half-life of 4 h (Fig. S1 and methods). The predicted *vs*. observed concentrations in the central and peripheral compartments of the HFIM fit well in all experiments reported thereafter, including those with the combination of ciprofloxacin and phages (Fig. 1). These data demonstrate the reproducibility of the disposition of ciprofloxacin in the HFIM and reveal that the presence of phages in the peripheral compartment does not affect it.

**Figure 1.**
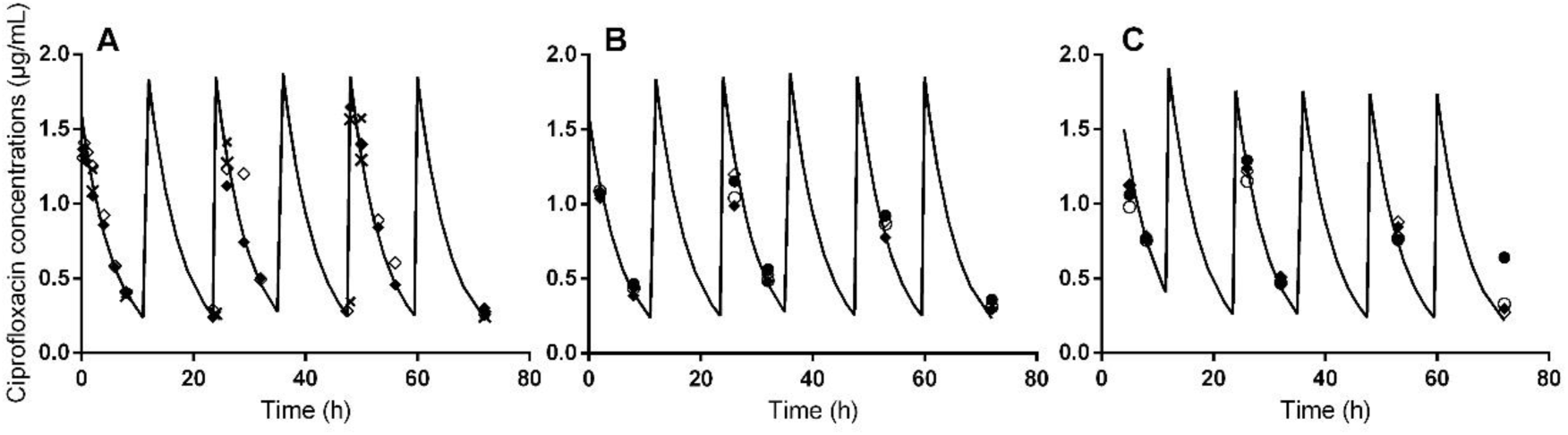
The regimen of ciprofloxacin administered in the HFIM reproduces the regimen of oral treatments. Expected (black line) and observed (diamonds and circles for standard and high inoculum, respectively) concentration-time profiles of ciprofloxacin in the HFIM (inoculated with *P. aeruginosa* strain PAK) after its administration twice a day for the following experiments: A, ciprofloxacin alone. B, ciprofloxacin administered simultaneously with the two-phage cocktail. C, ciprofloxacin administered 4 h post two-phage cocktail. n=2 for each inoculum represented by open and filled symbols. The concentrations in the peripheral compartment are represented with crosses in panel A.

### Clinically-relevant ciprofloxacin regimen selects rapidly for less susceptible clones

Following the inoculation of *P. aeruginosa* in the extracapillary space of the HFIM cartridge (Fig. S1 and S2), the bacterial concentration reached 5.7 log_10_ CFU/mL within 1 h, time point at which different treatments were administered. In the absence of treatment, this bacterial concentration increased to around 9.5 log_10_ CFU/mL over the first 24 h and remained roughly stable during the next 48 h (Fig. 2A).

**Figure 2.**
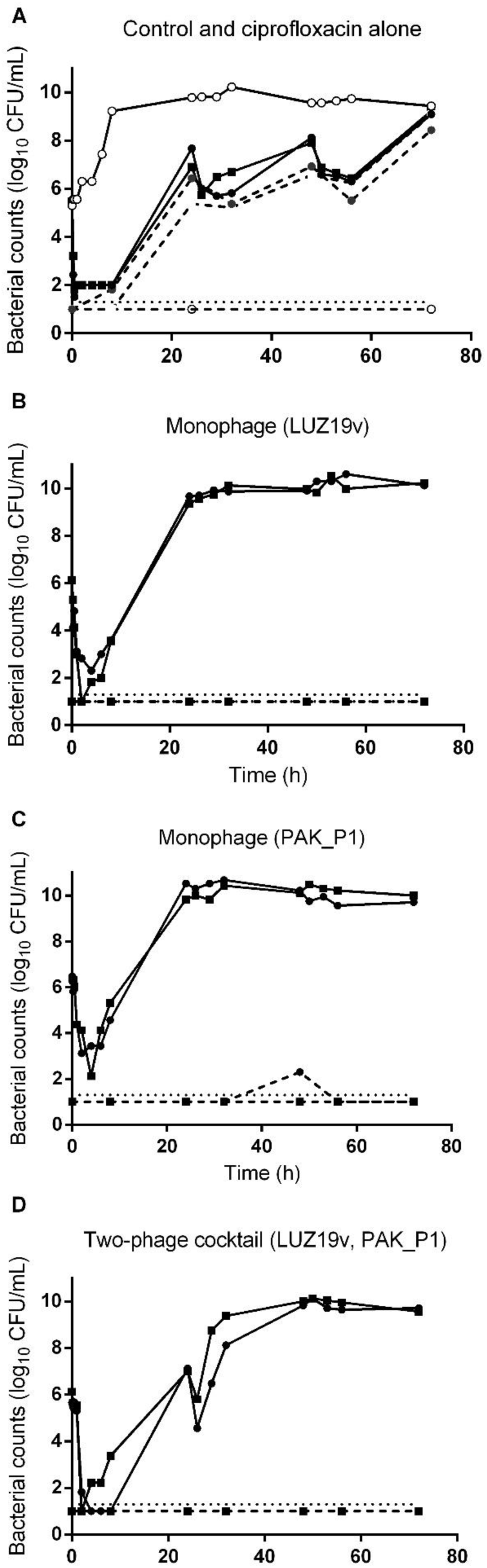
The growth of *P. aeruginosa* in the HFIM is not controlled by phages or ciprofloxacin. The concentration of *P. aeruginosa* strain PAK (log_10_ CFU/mL) in the HFIM from 1 h post-inoculation to72 h after exposure to ciprofloxacin or phages. A, control experiment (n=1) and ciprofloxacin alone (n=2). B, monophage (LUZ19v) (n=2). C, monophage (PAK_P1) (n=2). D, two-phage cocktail (LUZ19v, PAK_P1) (n=2). Solid lines represent total bacterial populations and dashed lines represent less-susceptible bacteria growing on agar containing 0.5 µg/mL of ciprofloxacin. Square and circles represent independent experiments. The horizontal dotted line corresponds to the LOD.

Within 30 min after the start of the ciprofloxacin regimen, the density of *P. aeruginosa* decreased by more than 3-logs and remained below the limit of detection (LOD) between 1 h and 8 h (Fig. 2A). Subsequently, the bacterial density increased, despite the ciprofloxacin regimen, reaching at 72 h a similar value (9.2±0.1 log_10_ CFU/mL) compared to the untreated control.

Samples from the HFIM were plated twice a day on agar supplemented with 0.5 µg/mL ciprofloxacin (8-fold MIC) to assess the selection of less-susceptible bacteria. No bacteria from the initial inocula (n=17) grew on this selective medium. In samples exposed to ciprofloxacin, the density of less-susceptible bacteria increased over the 72 h to reach 43-100% of the population (Fig. 2A). The MIC of ciprofloxacin for the bacteria sampled at 72 h increased by 250 fold (16 µg/mL; Table 1), showing that within 24 h, the ciprofloxacin regimen administered in the HFIM selected for less-susceptible bacteria.

**Table 1.** MIC of ciprofloxacin of the parental strain PAK and clones from samples exposed to either ciprofloxacin, or phages, or their combination in the HFIM during 72 h. Grey lines correspond to experiments performed with an inoculum of 8 log_10_ CFU/mL NBR, no bacteria recovered

| Experimental conditions | MIC ($\mu\text{g/mL}$) |
| --- | --- |
| Inoculum ($5.7 \log_{10} \text{CFU/mL}$) | 0.064 |
| **HFIM (72 h samples)** |  |
| Control (n=1) | 0.064 |
| Ciprofloxacin (n=2) | 16; 16 |
| LUZ19v (n=2) | 0.128; 0.128 |
| PAK_P1 (n=2) | 0.064; 0.064 |
| LUZ19v and PAK_P1 (n=2) | 0.064;0.064 |
| Simultaneous combination of ciprofloxacin and phages (n=2) | NBR |
| Delayed combination of ciprofloxacin and phages (n=2) | NBR |
| Simultaneous combination of ciprofloxacin and phages (n=2) | 1; 2 |
| Delayed combination of ciprofloxacin and phages (n=2) | NBR |

NBR, no bacteria recovered

### Single or two-phage local administration selects for phage-resistance

We next assessed with the HFIM the susceptibility of *P. aeruginosa* to two phages, the *Myoviridae* PAK_P1 and the *Podoviridae* LUZ19v, both positively evaluated previously for the treatment of acute lung infections in mice [20,21]. The frequency of bacteria among the naïve population that could grow in the presence of LUZ19v and PAK_P1 was 3×10^−7^ and 6×10^−5^, respectively (Table 2).

**Table 2.**
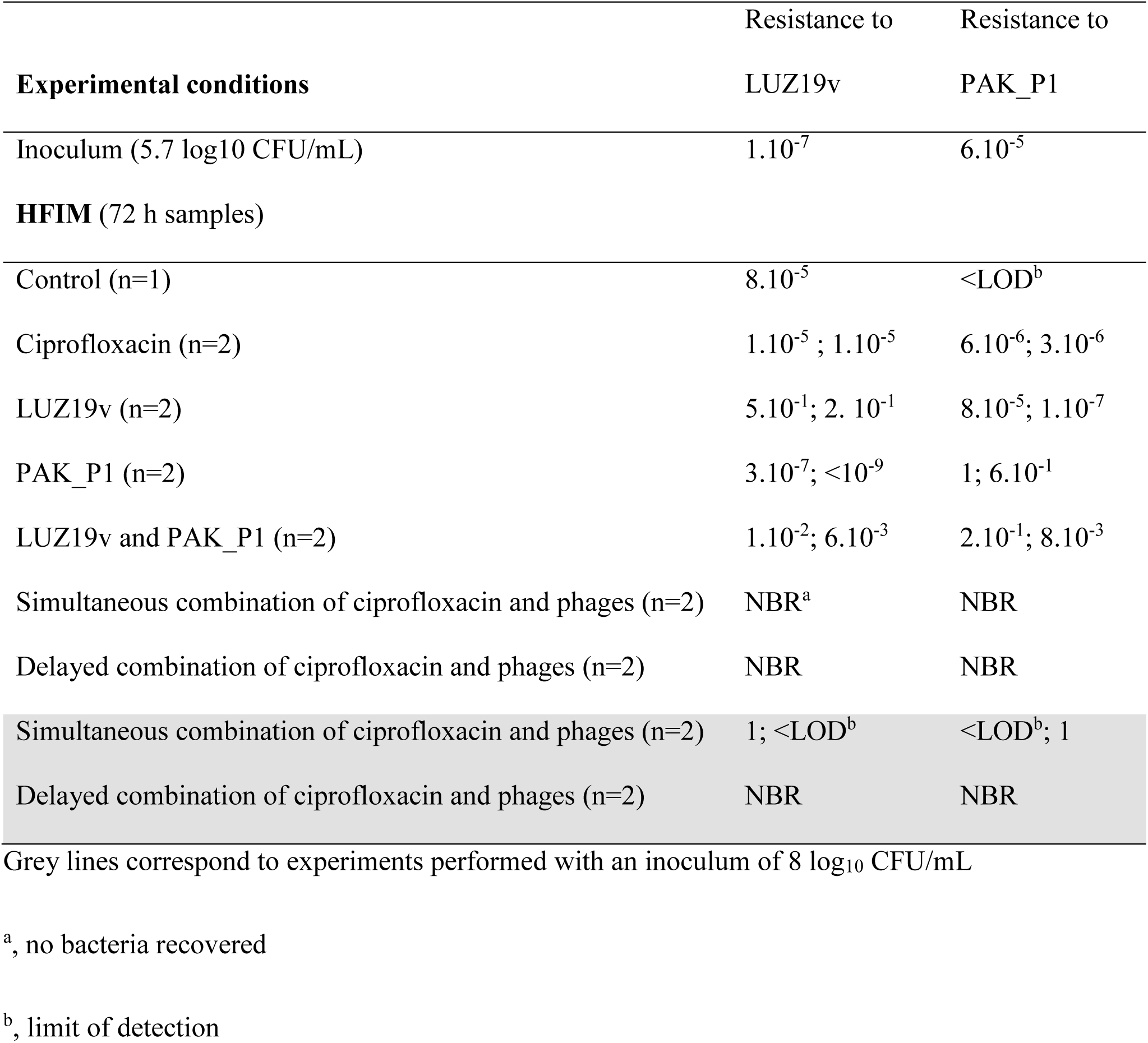
Frequencies of phage resistant clones from samples exposed to either ciprofloxacin, or phages, or their combination in the HFIM during 72 h Grey lines correspond to experiments performed with an inoculum of 8 log_10_ CFU/mL

Phages were administered once in the extracapillary space of the HFIM cartridge containing *P. aeruginosa* to mimic a local administration (Fig. S1). Along these experiments, no phage was detected in any of the samples taken from the central compartment, confirming that phages were strictly maintained in the extracapillary space (Fig. S2). The single dose of phages was set to obtain 7.5 log_10_PFU/mL in the HFIM (8.8 log_10_PFU *in toto*), which corresponds approximately to a phage:bacteria ratio of 100 at the time of administration.

Following the administration of phage LUZ19v or PAK_P1, the *P. aeruginosa* density dropped to 1.9±1.3 or 2.8±0.9 log_10_ CFU/mL within 2 h, or 4 h, respectively, and then started to increase continuously, reaching the density of the untreated control 24 h post-phage administration, and remained stable for another 48 h (Fig. 2B and 2C). Corresponding to the drop of bacteria, the density of LUZ19v or PAK_P1 increased during the first time points and reached a maximum at 24 h or 48 h, respectively (Fig. S3). The susceptibility to phages of samples taken at 72 h revealed that bacteria exposed to LUZ19v or PAK_P1 became nearly all resistant (20 to 50% and 60 to 100%, respectively), while keeping a large susceptibility to the second phage (<10^−5^ and 10^−7^, respectively) (Table 2). Therefore, monophage treatments were as inefficient as ciprofloxacin to control the growth of resistant bacteria in the HFIM during 72 h.

Since phages LUZ19v and PAK_P1 recognize two different bacterial receptors, the type IV pilus [22] and the lipolysaccharide (LPS, [18]), respectively, we assessed if their combination could lower the selection of resistant clones. The two-phage cocktail led to a similar reduction of the bacterial density during the first time points compared to phage LUZ19v alone (Fig. 2D). Then, the slope of the bacterial growth between 8 and 24 h was less steep compared to monophage treatments. Bacterial counts reached similar levels to the untreated control at 48 h and in the 72 h samples. The proportion of bacteria resistant to either phages was about one order of magnitude lower compared to single treatments (Table 2). Therefore, the use of two phages instead of one delays the selection of phage-resistant bacteria but does not prevent it.

The MIC of bacteria at 72 h following exposure to one or two phages was similar to the MIC of the inoculated strain showing that the exposure to phages does not select for less susceptible clones to ciprofloxacin (Table 1). Reciprocally, the bacteria from the HFIM exposed to ciprofloxacin were as susceptible to either phages as the control (Table 2).

### The combination of phages with ciprofloxacin prevents the growth resistant bacteria

To assess the impact of the combination of phages with ciprofloxacin, we tested two modalities corresponding to a simultaneous or a delayed treatment (phages first and ciprofloxacin 4 h later). The simultaneous treatment led to a rapid killing of bacteria, since their density dropped below the LOD in 15 min (Fig. 3A). Impressively, we could not detect any colony on samples taken during the next 72 h. During these experiments (n=2), the density of phages was stable, suggesting that they did not amplify (Fig. S4). When adding ciprofloxacin 4 h after the two-phage cocktail, the density of phages slightly increased and then remained stable up to 72 h. Here, again, the combination rapidly killed the population of *P. aeruginosa*, and no viable bacteria were recovered at any time after 1 h (Fig. 3B). Similar results were obtained when either phage was simultaneously added with ciprofloxacin (Fig. S5).

**Figure 3.**
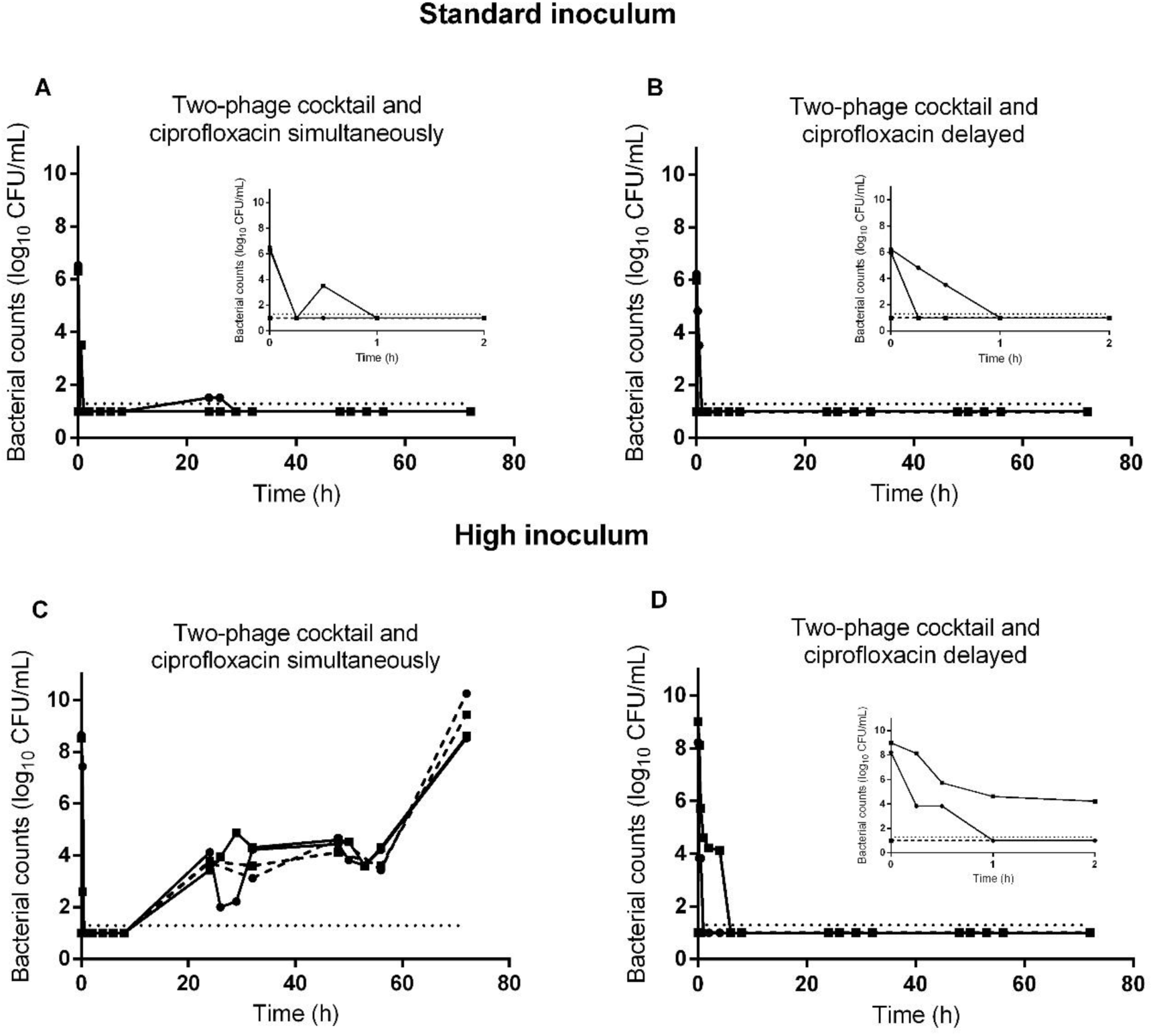
The growth of *P. aeruginosa* in the HFIM is only controlled by the combination of phages and ciprofloxacin. Population of bacteria (mean±SD in log_10_ CFU/mL) in the HFIM over 72 h post exposure of a standard (A and B) or high inoculum (C and D) of *P. aeruginosa* to the combination of ciprofloxacin and phages. A and C, combination of simultaneous administrations of ciprofloxacin with the two-phage cocktail (n=2). B and D, combination of the two-phage cocktail with ciprofloxacin administrated 4 h post-phages (n=2). Solid lines represent total bacterial populations and dashed lines represent less-susceptible bacteria growing on agar containing 0.5 µg/mL of ciprofloxacin. Square and circles represent independent experiments. The horizontal dotted line corresponds to the LOD.

### The antibacterial efficacy of the combination of phages with ciprofloxacin is density dependent

To assess the robustness of the combined treatment, the HFIM was inoculated with a 1000-fold higher bacterial inoculum while the regimen of either ciprofloxacin and phages remained unchanged. Following such inoculation, the bacterial density reached 8.6±0.3 log_10_ CFU/mL within 1 h, which corresponds to a phage:bacteria ratio of 0.1. The simultaneous addition of phages and ciprofloxacin rapidly killed bacteria with a density falling below the LOD between 30 min and 8 h (Fig. 3C). Subsequently, the bacterial density increased reaching 8.6±0.06 log_10_ CFU/mL at 72 h. The bacteria recovered had a reduced susceptibility to ciprofloxacin (16 to 32-fold higher MIC) and to phages compared to the naïve population (Tables 1 and 2). In the two independent assays, the proportion of bacteria resistant to each phage increased with bacteria becoming fully resistant to LUZ19v and partially resistant to PAK_P1 in one replicate and the other way around in the second.

By contrast, when adding ciprofloxacin 4 h after the two-phage cocktail, the initial reduction of bacteria was slower compared to the simultaneous administration with bacterial density falling below the LOD at 1 h for one replicate and 6 h for the other (Fig. 3D). However, after this decline, no increase of the bacterial density was observed and again no colony could be recovered on samples taken during the next 72 h. We concluded that when phages reduce first the size of the bacterial population, the remaining population was not large enough to include less-susceptible mutants to ciprofloxacin.

## Discussion

*P. aeruginosa* lung infections are increasingly difficult to treat with antibiotics calling for therapeutic strategies to enhance bacterial killing. In this study, we developed an innovative use of the HFIM to evaluate the potential benefit of combining ciprofloxacin with phages. The HFIM allows the simulation of a clinically relevant ciprofloxacin concentration profile obtained with oral treatment that leads in monotherapy to the selection of clones with increasing resistance as previously reported [9–11]. The combination of ciprofloxacin and meropenem in the HFIM suppressed the growth of resistant *P. aeruginosa* isolates, including hypermutable strains [10,11]. In clinics, this combination requires IV administration and the hospitalization of patients [2]. Moreover, it participates to the upscaling of drugs use and overall increases the selection of MDR strains.

Instead of a second antibiotic, we evaluated the combination of ciprofloxacin with phages. A unique local administration of one or two phages could not prevent the growth of phage-resistant bacteria, as generally observed with *in vitro* tests. We observed that the growth of resistant clones in presence of the two phages was delayed compared to single phages, in agreement with increased fitness cost, as reported elsewhere [23,24].

In contrast to previous studies that combined a single addition of ciprofloxacin, with either phage OKMO1, or PEV31, or a five-phages cocktail, we did not observe a modification of the MIC for ciprofloxacin in any of the phage-resistant clones tested [17,25,26]. This suggests that these clones may not be selected when a clinically relevant regimen of ciprofloxacin is used. Moreover, the ciprofloxacin resistant clones remained largely susceptible to each of the two phages as their frequency was the same as in the control (without treatment). Altogether, the lack of correlation between the profiles of susceptibility and resistance to ciprofloxacin and the two phages demonstrates their independent antibacterial activity.

The combination of ciprofloxacin with one or two phages administered simultaneously on the HFIM inoculated with the same bacterial density than individual treatments abolished the growth of resistant clones during at least 72 h, a long-term performance compared to less than 30 h for the latter treatments.This strongly suggests that in these conditions no bacteria survived. However, when the initial bacterial density was 1000 fold higher the growth of resistant clones was detected at 24 h and rise up during the next 48 h, showing that this regimen was unable to control a dense population of *P. aeruginosa*. When phages are administered first and the ciprofloxacin 4 h later, the drop of bacteria aligned with the increase of phage concentrations. The bacterial density reached the LOD until the end of each experiment, with low or high inoculum, suggesting that no bacteria survived these regimens.

The *in vivo* efficacy of the combination of ciprofloxacin (a single oral dose simulating 750 mg in human) with phages (a single intravenous administration of 10^10^ PFU) was previously tested in an experimental *P. aeruginosa* endocarditis in rats and led to more frequent negative vegetation cultures after 6 h than in rats receiving only phages or only ciprofloxacin [27]. Using a murine model of *P. aeruginosa* pulmonary infection, a unique dry power insufflation of ciprofloxacin and phages led to a reduction of nearly 6 log_10_ CFU in 24 h [28]. In these two studies, a unique dose of ciprofloxacin was used associated to short time end-points. The data we obtained with the HFIM suggest that the administration of antibiotics following their recommended regimens could increase the efficacy of these combinations on longer time points.

One of the limitations of our study relates to the lack of an immune component that could enhance the overall efficacy of such regimen as the immune system was previously shown to cooperate with phages during experimental pulmonary phage therapy [29]. Another limitation is the lack of loss of phages over time as they remained trapped in the same compartment of bacteria. However, the decay of phages in uninfected or infected lungs of mice was shown to be rather weak (below 1-log per day) compared to the overall density of phages in the HFIM [30]. The data presented here advocate in favor of a translation to clinics that could ultimately slow down the use of multiple antibiotics and therefore, the selection of MDR strains [17].

## Acknowledgements

We thank Dwayne Roach for the *in vitro* adaptation of LUZ19v to strain PAK, and Thierry Pédron for the preparation of phage lysates.

This project was supported by a grant from Institut Carnot France Futur Elevage and Institut Carnot Pasteur Maladies Infectieuses to AB and LD, respectively.

## Supplementary figures

**Figure S1.**
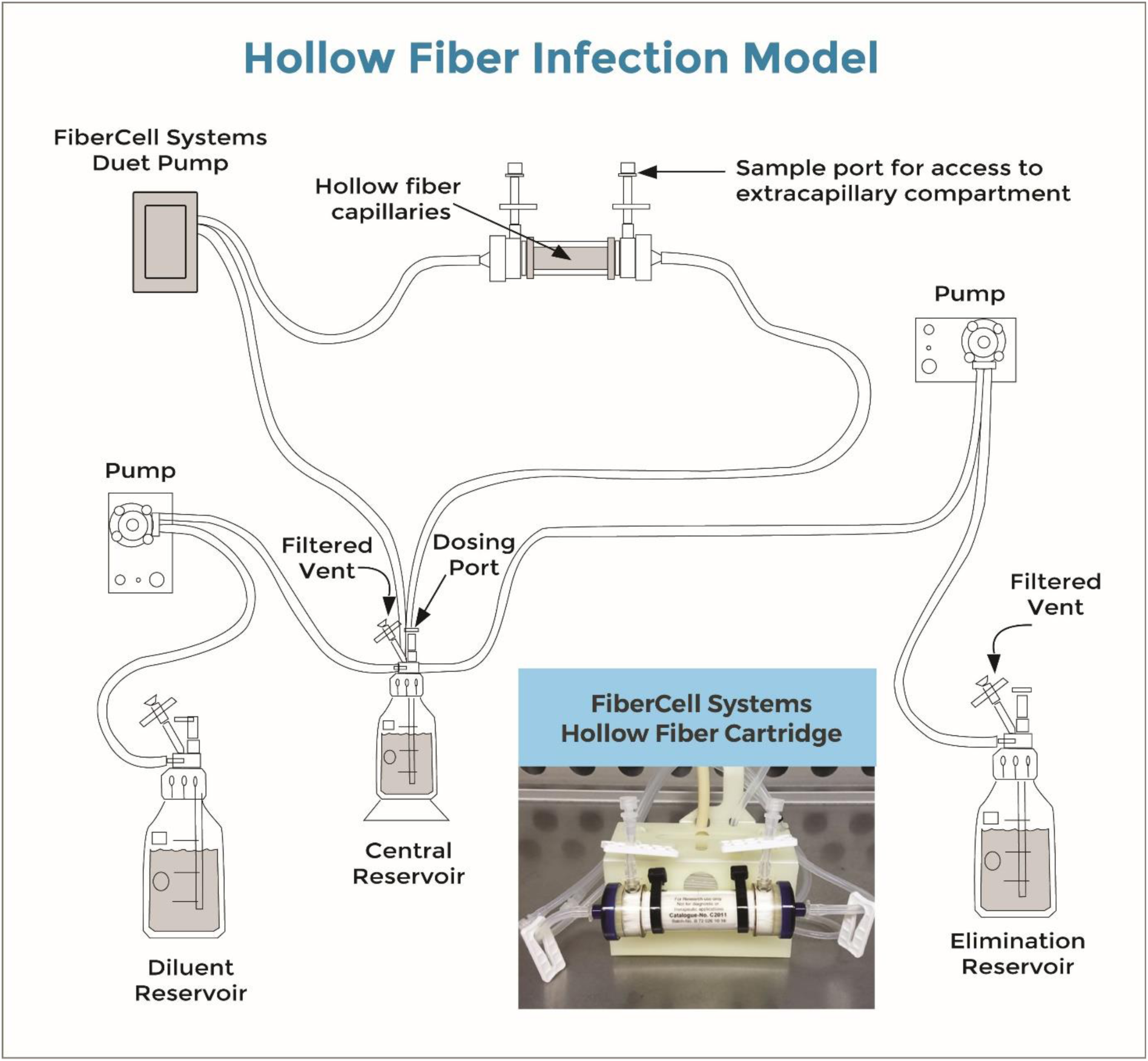
Schematic representation of the Hollow Fiber Infection Model kindly provided by FiberCell Systems®. Bacteria and phages were trapped in the extracapillary space of the cartridge (peripheral compartment) (see also embedded photo). Ciprofloxacin was added to the central reservoir and freely circulated through the cartridge and bacteria by means of the Fibercell Systems Duet pump® (FiberCell Systems, Inc., Frederick, MD, USA). Ciprofloxacin concentrations decreased over time after drug administrations, due to the continuous addition of a diluent (MHB) by means of another set of pumps.

**Figure S2.**
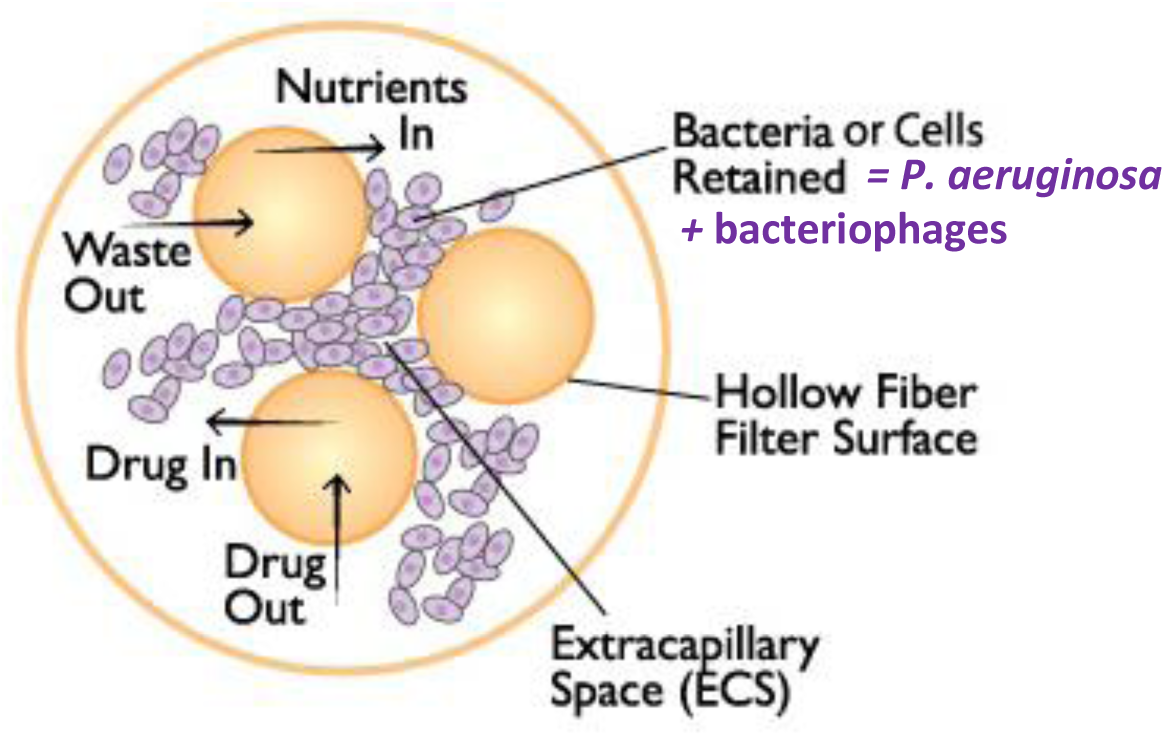
Schematic representation of the cross-section of the cartridge of the HFIM, kindly provided by FiberCell Systems®. Phages were directly introduced in the extracapillary space containing *P. aeruginosa* to simulate local administration. Both phages and *P. aeruginosa* remained trapped in this extracapillary space during the experiments. The drug, here ciprofloxacin, freely circulates through the fibers and was distributed both in the central reservoir and the extracapillary spaces of the cartridge.

**Figure S3.**
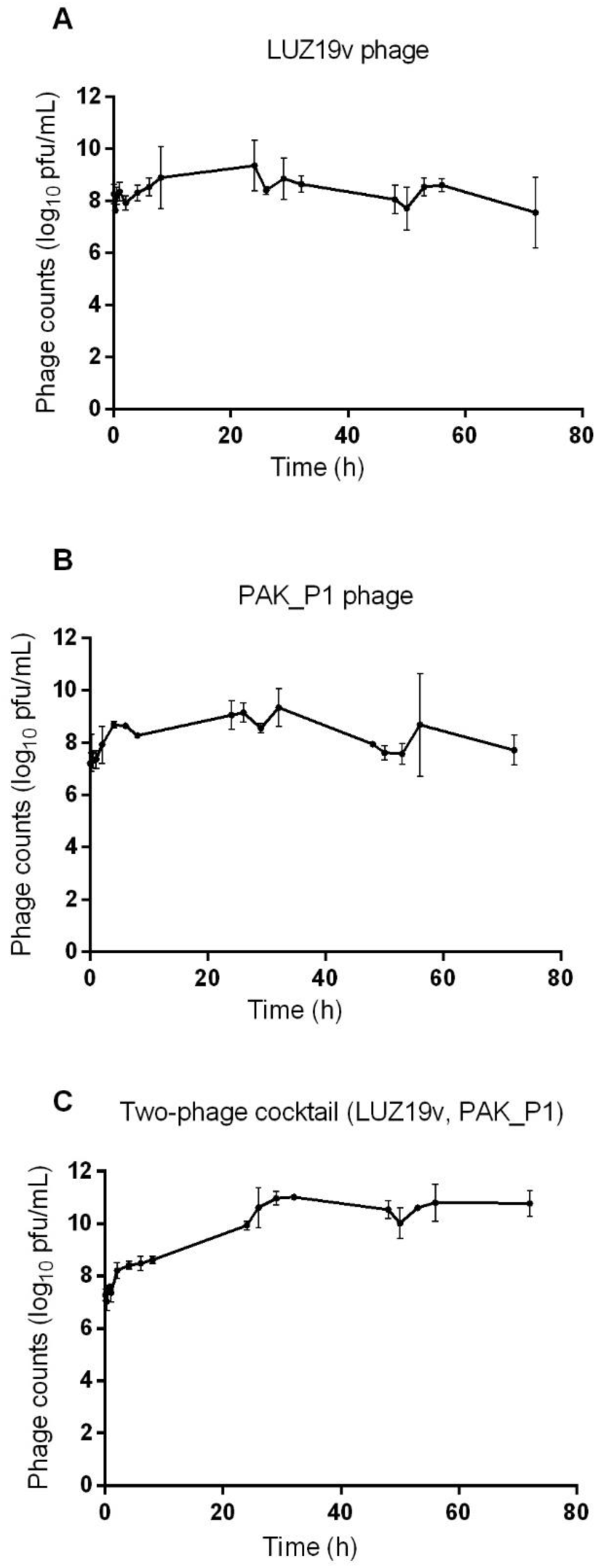
Total population of phages (mean±SD in log_10_ PFU/mL) in the HFIM over 72 h in the different experiments. A, phage LUZ19v alone (n=2). B, phage PAK_P1 alone (n=2). C, phages LUZ19v and PAK_P1 (n=2).

**Figure S4.**
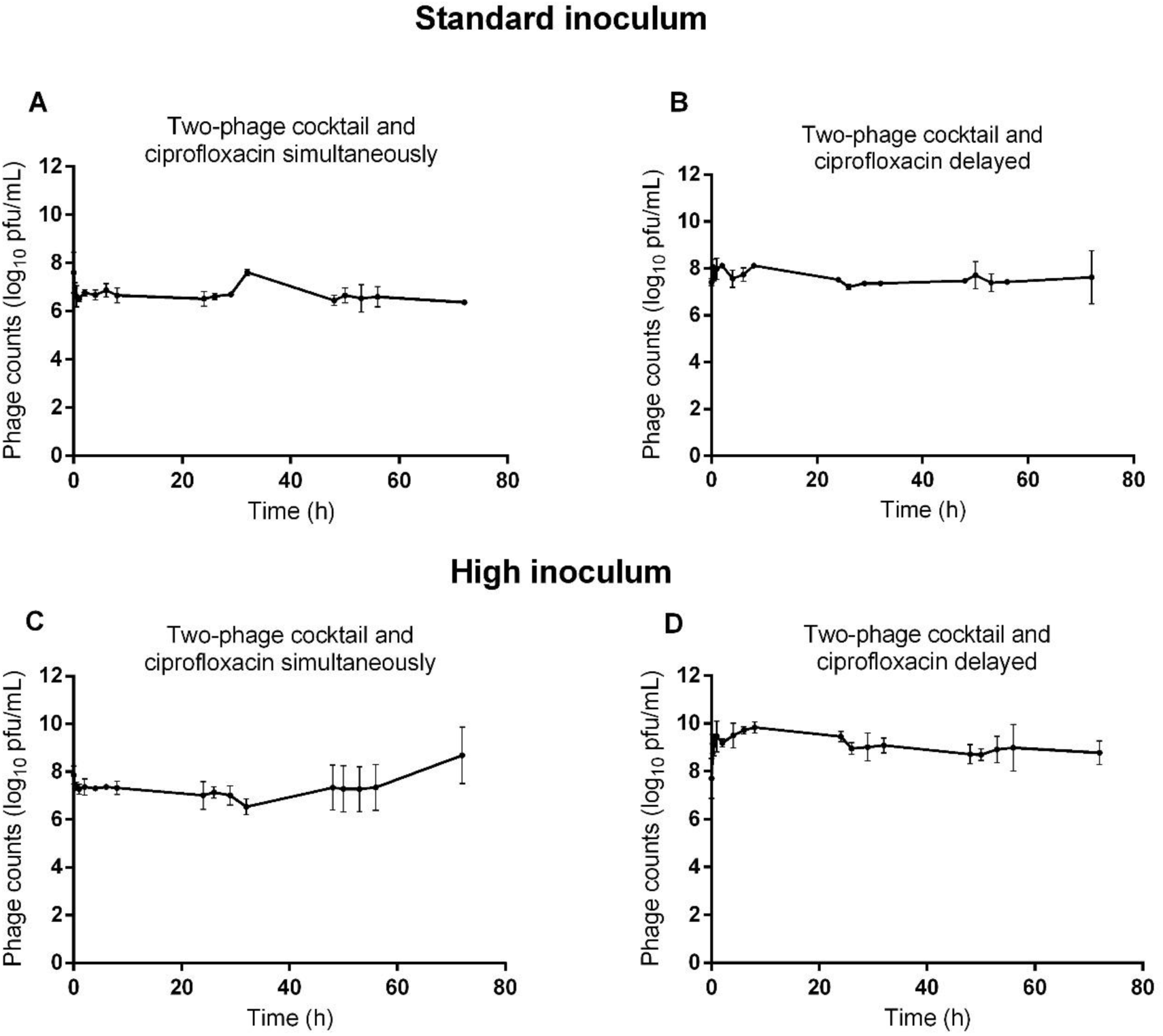
Total population of phages (mean±SD in log_10_ PFU/mL) in the HFIM over 72 h in the experiments with a standard (A and B) or high inoculum (C and D) of *P. aeruginosa*. A and C, combination of simultaneous administrations of ciprofloxacin with the two-phage cocktail (n=2). B and D, combination of the two-phage cocktail with ciprofloxacin administrated 4 h post-phages (n=2).

**Figure S5.**
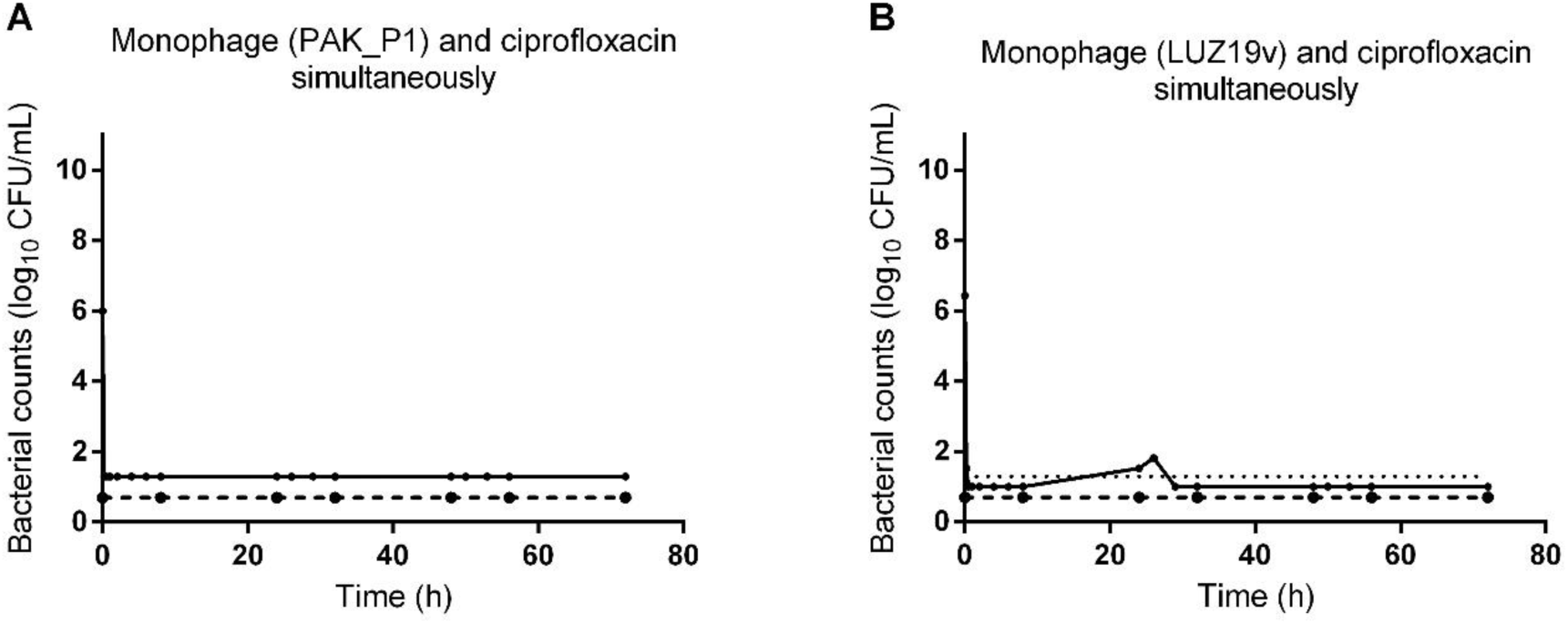
Population of bacteria (mean±SD in log_10_ CFU/mL) in the HFIM over 72 h after exposure to the combination of ciprofloxacin and a single phage. A, combination of simultaneous administration of ciprofloxacin with LUZ19v (n=1). B, combination of simultaneous administration of ciprofloxacin with PAK_P1 (n=1). Solid lines represent total bacterial populations and dashed lines represent less-susceptible bacteria growing on agar containing 0.5 µg/mL of ciprofloxacin. The horizontal dotted line corresponds to the LOD.

## Supplementary file

Samples for ciprofloxacin quantification were withdrawn from the central compartment at 0.25, 0.5, 1, 2, 4, 6, 8, 24, 26, 29, 32, 48, 50, 53, 56, and 72 h, and from the extracapillary space of the cartridge at 2, 8, 24, 26, 48, 54, and 72 h (Supplementary file). Samples were centrifuged at 3000 g for 10 min, and the supernatant was stored at -20°C. One hundred microliters of water containing the marbofloxacin internal standard at 5 µg/mL were added to 100 µL of calibrators, quality controls, or samples. The mixture was vortexed at 1400 rpm for 2 min at 10°C and centrifuged at 20000 g for 10 min. The supernatant (20 µL) was injected into an Acquity ultra performance liquid chromatography (UPLC®) coupled to a UV detector (Waters, Milford, MA, USA). Ciprofloxacin was eluted at 0.3 mL/min on an Acquity UPLC BEH C18 column (2.1×50 mm, 1.7 µm) equipped with a frit (0.2 µm, 2.1 mm) and set at 40°C under the following gradient conditions: t0 90% A (H_2_O acidified with 0.1% HCOOH) 10% B (acetonitrile); t(4 min) 60% A and 40% B. The return to initial conditions was held for 1 min. Wavelength detection was set at 278 nm. Chromatographic data were monitored by Empower software (Waters, Milford, MA, USA). The method was validated from 0.05 to 5 µg/mL of ciprofloxacin with a linear model weighted by 1/X^²^ (X=concentration). Precisions and accuracy were checked by injecting six replicates of QC samples over three days, at the limit of quantification (0.05 µg/mL); 0.075 µg/mL; 0.75 µg/mL, and 4 µg/mL. Accuracies ranged from 92% to 107%, with intra-day and inter-day CV precisions below 5% and 13%, respectively. The limit of quantification was validated at 0.05 µg/mL, with an accuracy of 96% and intra-and inter-day CV precision lower than 6%.

